# Apparent size and morphology of bacterial microcompartments varies with technique

**DOI:** 10.1101/864074

**Authors:** Nolan W. Kennedy, Jasmine M. Hershewe, Taylor M. Nichols, Eric W. Roth, Charlene D. Wilke, Michael C. Jewett, Danielle Tullman-Ercek

## Abstract

Bacterial microcompartments (MCPs) are protein-based organelles which encapsulate metabolic pathways. Metabolic engineers have recently sought to repurpose MCPs to encapsulate heterologous pathways to increase flux through pathways of interest. As MCP engineering becomes more common, standardized methods for analyzing changes to MCPs and interpreting results across studies will become increasingly important. In this study, we demonstrate that different imaging techniques yield variations in the apparent size of purified MCPs from *Salmonella enterica* serovar Typhimurium LT2, likely due to variations in sample preparation methods. We provide guidelines for preparing samples for MCP imaging and outline expected variations in apparent size and morphology between methods. With this report we aim to establish an aid for comparing results across studies.

## Introduction

Scientific research has recently come under fire for what is being dubbed a crisis of reproducibility. Current studies estimate that 75-90% of findings in high-profile journals are not reproducible [1]. The issue has seeped into fields across every domain of scientific inquiry [2]. While the cause of any given irreproducible result will vary from case to case, a lack of technique standardization across studies can lead to artefactual results or false conclusions [3]. In fields in which different techniques are employed to test similar hypotheses, it is important to place results into the proper context and understand the limitations of each technique. Here, we provide guidelines for technique standardization and result interpretation in the bacterial microcompartment engineering field.

Bacterial microcompartments (MCPs) are protein-based organelles found in diverse species of bacteria [4–6]. These were originally identified in cyanobacteria and were hypothesized to be viruses based on their appearance [7,8]. However, these structures were later determined to be important for the carbon concentrating mechanism for certain species of autotrophic microbes [9–12]. Since then, numerous diverse types of MCPs have been identified in species ranging from cyanobacteria and halophilic ocean-dwelling bacteria, to enteric pathogens and soil-dwelling microbes [13,14]. In addition to the cyanobacterial compartments used for carbon fixation, many MCPs are used by enteric pathogens for the metabolism of unique carbon sources that move through toxic or volatile intermediates, imparting a competitive advantage [15–18].

The flagship archetype for metabolic MCPs is the 1,2-propanediol utilization (Pdu) MCP found in *Salmonella enterica.* The Pdu MCP encapsulates the enzymatic machinery necessary for metabolism of 1,2-propanediol (1,2-PD), a carbon source found in the gut of *Salmonella* hosts [15]. The 1,2-PD metabolic enzymes are surrounded by a protein shell composed of multiple types of trimeric, pentameric, and hexameric shell proteins. The reported size of these irregularly-shaped protein organelles varies widely from 77-220 nm in diameter (**S1 Table**), and rigorous methods for size quantification are sparse [8,13,15,19–23].

The Pdu MCP has been studied in-depth since the early 1990s, but it has recently increased in popularity due to its potential utility in metabolic engineering [24–26]. Metabolic engineers have sought to increase flux through target pathways of interest by increasing local concentrations of enzymes and their substrates [27]. MCPs can accomplish this task and offer the potential added benefit of sequestering toxic or volatile intermediates from damaging or escaping the cell [28,29]. They also have the potential to reduce unwanted side reactions and provide private cofactor pools separate from central metabolism [30].

Recent efforts to engineer MCPs focused on loading heterologous proteins to the lumen of these structures, as well as modifying the MCP shell to alter substrate and product diffusion [23,31]. Even modest engineering efforts can affect the size, shape, and morphology of MCPs. For example, knocking out or over-expressing different shell proteins leads to dramatic changes in the shape of MCPs, with many appearing to be long, hollow tubes [32–36]. As engineering efforts continue, it will become increasingly important to have a standardized set of tools for the field to determine and compare the size, shape, and morphology of engineered or altered MCPs across different studies. To date, there is no widely agreed-upon method for visualizing and measuring MCPs, with labs across the field utilizing their own preferred technique. Here we demonstrate that different techniques can yield variable apparent results, even on identical samples. We provide an outline for choosing an appropriate technique and subsequently correlate the results across the many visualization and sizing techniques used in the field.

## Methods

### Microcompartment Expression and Purification

Intact Pdu MCPs were purified from lysed cultures of *Salmonella enterica* serovar Typhimurium LT2 using a centrifugation process as previously described [37–39]. Briefly, starter cultures were grown in 5 mL of LB-Miller for 24 hours at 30°C, 225 RPM and subsequently subcultured 1:1000 into 200 mL of no carbon-E (NCE) minimal media (29 mM KH_2_PO_4_, 34 mM K_2_HPO_4_, 17 mM Na(NH_4_)HPO_4_, 1 mM MgSO_4_, and 50 μM ferric citrate) supplemented with 42 mM succinate as a carbon source and 55 mM 1,2-propanediol for MCP induction. NCE cultures were grown at 37°C, 225 RPM to a final target OD_600_ of ∼1-1.5, after which they were harvested and lysed. The lysed cultures were centrifuged at 12,000 x G for 5 minutes to remove cell debris. MCPs were then pelleted from the resulting supernatant through centrifugation at 21,000 x G for 20 minutes and collected. The total protein concentration of MCP samples was measured using the Pierce™ BCA Protein Assay Kit (Thermo Scientific) per the manufacturer’s instructions, and concentrations were normalized as necessary for each analysis method. All MCP samples were stored at 4°C until use and were prepared for analysis within 5 days of purification to avoid MCP aggregation and degradation [40].

### Protein Electrophoresis

Purified MCPs were assessed for composition by SDS-PAGE as previously described [38]. Briefly, MCP samples were boiled in Laemmli buffer at 95°C for 5-7 minutes. The denatured samples were then loaded onto 15% SDS-PAGE gels and separated at 120 V for 90 minutes. Approximately 2-2.25 µg total protein was loaded for each sample, as measured by BCA assay. Gels were then stained with Coomassie and imaged using the Bio-Rad ChemiDoc XRS+ (**S1 Fig**).

### Negative-Stain Transmission Electron Microscopy

Samples were set on 400 mesh Formvar-coated copper grids (EMS Cat# FF400-Cu) with a carbon film. Grids were treated by glow discharge using a PELCO easiGlow glow discharge cleaning system for a total of 10 seconds at 15 mA. MCP samples were placed onto the grids immediately following glow discharge. We found that staining and contrast were best if MCPs were left undiluted (between 0.5-1.0 mg/mL). A volume of 10 μL of purified MCPs was pipetted onto the surface of the grids, which were held in place by negative-action tweezers. The samples were allowed to sit for 2 minutes before being wicked away with filter paper. Note that some of the liquid should always be left on the grid to avoid sample collapse. The samples were washed three times by dipping the grid in a small droplet of deionized water for three seconds. The samples were fixed by placing 10 μL of 2% (v/v) glutaraldehyde onto the grid for 2 minutes. Note that glutaraldehyde should be stored under N_2_, and the 2% dilution should be made fresh before each sample preparation session. After the 2-minute incubation, the glutaraldehyde was wicked away using filter paper and the sample was washed three times in deionized water. Samples were stained with 1% (w/v) aqueous uranyl acetate (UA) by applying 10 μL of UA to the grids for 2 minutes. The UA was wicked away completely using filter paper. Note that all samples, fixative, stain, and deionized water were spun at 12,000 x G for 2 minutes before use to remove any aggregates. Samples were imaged at the Northwestern Electron Probe Instrumentation Center (EPIC) using the Hitachi HT-7700 Biological S/TEM Microscope and the Galtan Orius 4k × 2.67k digital camera.

For samples that were exchanged into solvent to prevent collapsing, samples were first fixed as described above in 2% glutaraldehyde. The samples were exchanged into 30% (v/v) ethanol for 1 minute, then 50% (v/v), 70% (v/v), and 90% (v/v) ethanol, followed by 100% ethanol three times. After this exchange into ethanol, samples were exchanged into 50% (v/v) and then 100% hexamethyldisilazane (HMDS). Samples exchanged into 100% HMDS were stained with UA as described above.

### Scanning Electron Microscopy

Samples were spotted and fixed onto 400 mesh Formvar-coated copper grids (EMS Cat# FF400-Cu) and processed through a 100% ethanol exchange as described above. Grids were placed into a sample holder and loaded into a Tousimis critical point dryer. The critical point dryer was run for a 10-minute purge cycle. Grids were mounted onto SEM stubs with carbon tape and coated with 6 nm of gold/palladium in a Cressington 208H sputter coater. Grids were imaged using a Hitachi SEM with 2 kV accelerating voltage and a 4 mm working distance.

### Cryo Transmission Electron Microscopy

Lacey Carbon 200 mesh Cu grids (EMS Cat# LC200-CU) were glow discharged in a Pelco easiGlow glow discharger for 30 seconds at 30 mA. 4 μL of sample solution was carefully pipetted onto the grids and plunge frozen in liquid ethane in a FEI Vitrobot Mark III with a blotting time of 5 seconds and blot offset of 0.5 mm. Grids were stored in liquid nitrogen and loaded into a Gatan 626.6 Cryo Transfer Holder cooled down to −170°C prior to observation in a JEOL JEM-1230 LaB6 emission TEM running at 100 kV. Images were collected with a Gatan Orius SC1000 CCD Camera, Model 831.

### Transmission Electron Microscopy of Ultra-thin Sections

Samples were pelleted at 21,000 x G in an Eppendorf 5424 microcentrifuge for 10 minutes. Pelleted samples were fixed with 2.5% (v/v) glutaraldehyde and 2% (v/v) paraformaldehyde in 0.1M PBS, post-fixed with 1% (w/v) OsO4 and 1% (w/v) UA, dehydrated in a graded series of ethanol, infiltrated with EMBed 812 epoxy resin, and embedded in beam capsules. The embedded samples were polymerized at 60°C for 48 hours prior to ultra-thin sectioning utilizing a Leica UC7 ultramicrotome. Sections were collected on 150 mesh Cu grids with a formvar/carbon membrane and stained with 3% (w/v) UA in 50% (v/v) methanol and Reynold’s lead citrate to further enhance contrast. The samples were observed in a JEOL JEM-1230 LaB6 emission TEM at 100 kV. Images were collected with a Gatan Orius SC1000 CCD Camera, Model 831.

### Dynamic Light Scattering Measurements

Samples were centrifuged at 12,000 x G for 5 minutes at 4°C immediately before dynamic light scattering (DLS) analysis to remove aggregated or insoluble protein. Dynamic light scattering was performed on a Zetasizer Nano ZS (Malvern Instruments Ltd., UK) with a measurement angle of 173°. Measurements were collected in triplicate at 4°C for 13 scans per measurement. Refractive index and temperature-adjusted viscosity were obtained from the instrument’s parameter library.

Nanoparticle tracking analysis was performed on a Nanosight NS300 using a 488 nm (blue) laser (Malvern Instruments Ltd., UK). Instrument settings were adjusted according to manufacturer recommendations. Measurements were collected for a duration of 60 s in 5 runs using a 1 mL syringe and a syringe pump speed of 30. Measurements were collected at room temperature.

### Image Analysis and Sizing

Images were contrast-adjusted and cropped using ImageJ [41]. For MCP sizing, images were scale-corrected based on the instrument used to collect the images. The oval tool was used to manually trace an ellipse surrounding MCPs. The longest diameter in the ellipse, corresponding to the widest diameter for the MCP, was recorded. Further data analysis was carried out using Microsoft Excel or R. Two-tailed t-tests were used to determine significance of differences between populations.

## Results and Discussion

### Negative-stain TEM of purified MCPs yields MCPs that appear deflated

Imaging MCPs using negative-stain transmission electron microscopy (TEM) is a standard technique used by the MCP field that has been widely adopted since Sinha and colleagues first described a method for MCP purification [39]. This technique enables clear identification of the border of each MCP, facilitating descriptions of shape and morphology (**Table 1**). Additionally, these results are generally higher contrast than techniques that involve imaging unpurified MCPs, such as TEM of ultra-thin cell sections.

**Table 1.** Comparison of different techniques used for MCP analysis. Table 1 lists the various techniques, along with their strengths and weakness, that are utilized in the MCP field and are assessed in this work. We have also included a brief list of specialized equipment necessary for each technique, and a list of previous works in the MCP field in which each technique was used. Our hope is that this will enable selection of the technique best-suited for each study.

| Method | Strengths | Weaknesses | Specialized Equipment | Previous Works |
| --- | --- | --- | --- | --- |
| **Transmission Electron Microscopy (TEM)** | •Relatively simple instrumentation, compared to other techniques<br>•Sample preparation is fast and straightforward<br>•Easy to see surface and shape morphology<br>•History of use in the field | •Compartments appear collapsed due to sample preparation methods<br>•Size analysis is slow compared to DLS, etc. | •Glow Discharge System<br>•Transmission Electron Microscope | [20,22–24,30,32,37,40,42–52] |
| **Ultra-thin section Transmission Electron Microscopy (TEM)** | •Can visualize compartments in native context (does not require purification)<br>•History of use in the field | •Relatively slow and difficult sample preparation, requiring multiple pieces of specialized equipment<br>•Cannot readily visualize surface morphology<br>•*In vivo* images are difficult to analyze due to other cellular components<br>•Due to the irregular shape of compartments, size determination using this method yields a wide distribution of apparent compartment diameters, skewing results | •Ultramicrotome<br>•Transmission Electron Microscope | [8,15,16,19,22,24,30,33–36,40,42–45,51,53–55] |
| **Scanning Electron Microscopy (SEM)** | •Compartments appear more true-to-size (less collapsed)<br>•Can visualize surface and shape morphology | •Compartments appear slightly collapsed<br>•Staining with a metal coat can hide surface morphologies and alter apparent size<br>•Sample preparation and imaging is relatively complex and requires multiple pieces of specialized equipment | •Glow Discharge System<br>•Critical Point Dryer<br>•Sputter Coater<br>•Scanning Electron Microscope |  |
| **Transmission Electron Cryo-microscopy (Cryo TEM)** | •Compartments retain solution size, shape, and morphology the best | •Sample preparation and imaging is difficult<br>•Contrast is low due to lack of staining | •Glow Discharge System<br>•Plunge freezer<br>•Cryo transfer holder<br>•Transmission electron microscope | [21,56] |
| **Dynamic Light Scattering (DLS), etc.** | •The most rapid, high-throughput method for determining the size distribution of a population of compartments | •Does not provide information on morphology | •DLS, Zeta Sizer, other system | [40,47,51] |

A drawback to the negative-stain TEM technique is that it requires sample dehydration as part of the sample preparation process. This ultimately leads to MCP collapse or deflation, as indicated in **Fig 1A**. Dark staining is present at the MCP interior, indicating that the stain is pooling in the collapsed, cup-shaped MCPs. To avoid this, fixing with glutaraldehyde is often used, but does not seem to completely prevent MCP collapse. MCPs appeared to be 102 ± 17 nm (mean ± standard deviation) in diameter when measured in images generated with this method (**Fig 2**).

**Fig 1.**
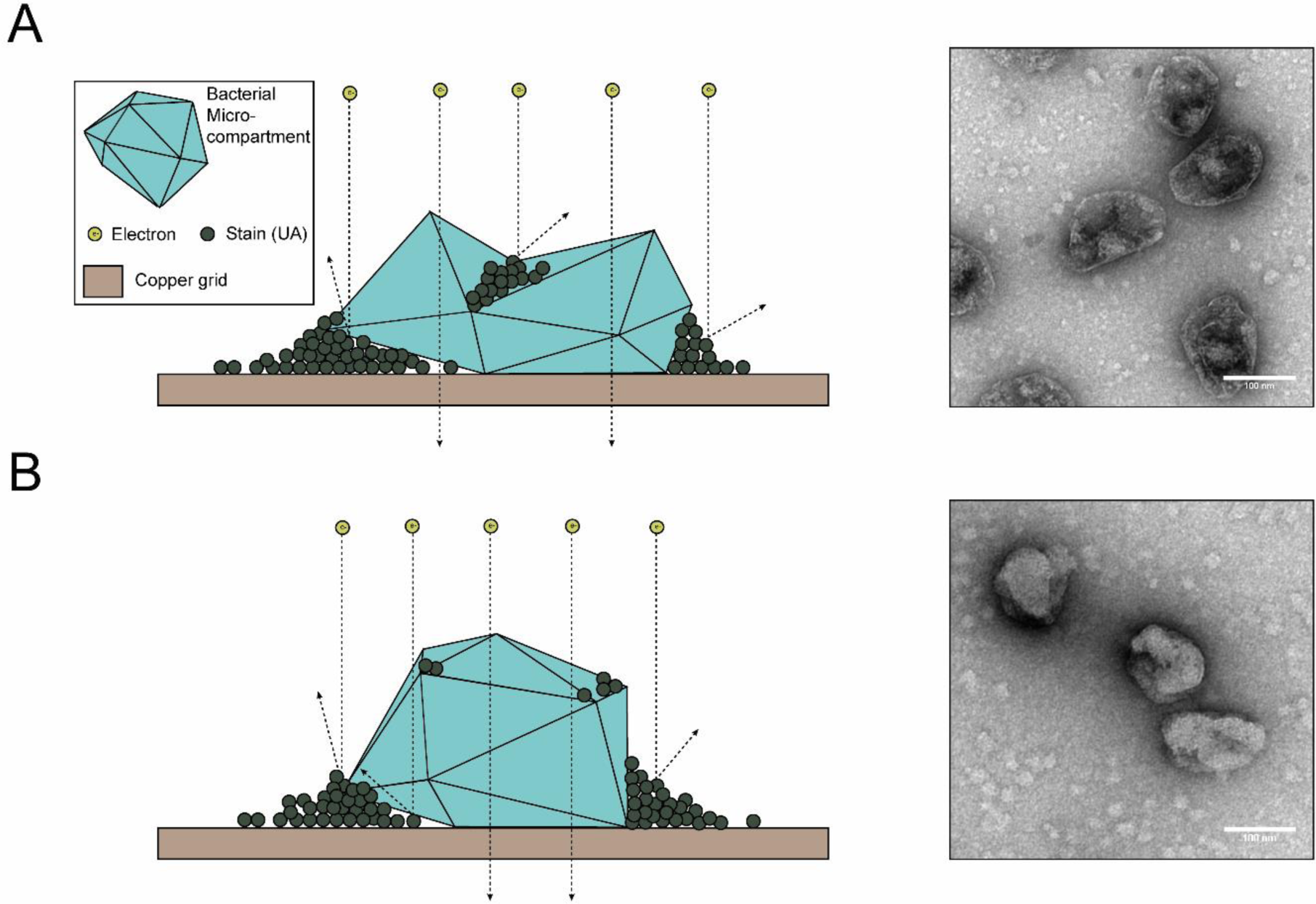
TEM of purified MCPs. (A) Schematic representation and transmission electron micrograph of negatively stained purified MCPs. Note that MCPs appear collapsed as evidenced by the pooled stain near the center of MCPs. (B) Schematic representation and transmission electron micrograph of negatively stained purified MCPs which were first dehydrated in ethanol and a high vapor pressure solvent (HMDS). Note that MCPs appear less collapsed than in (A). Scale bar (white) is 100 nm.

**Fig 2.**
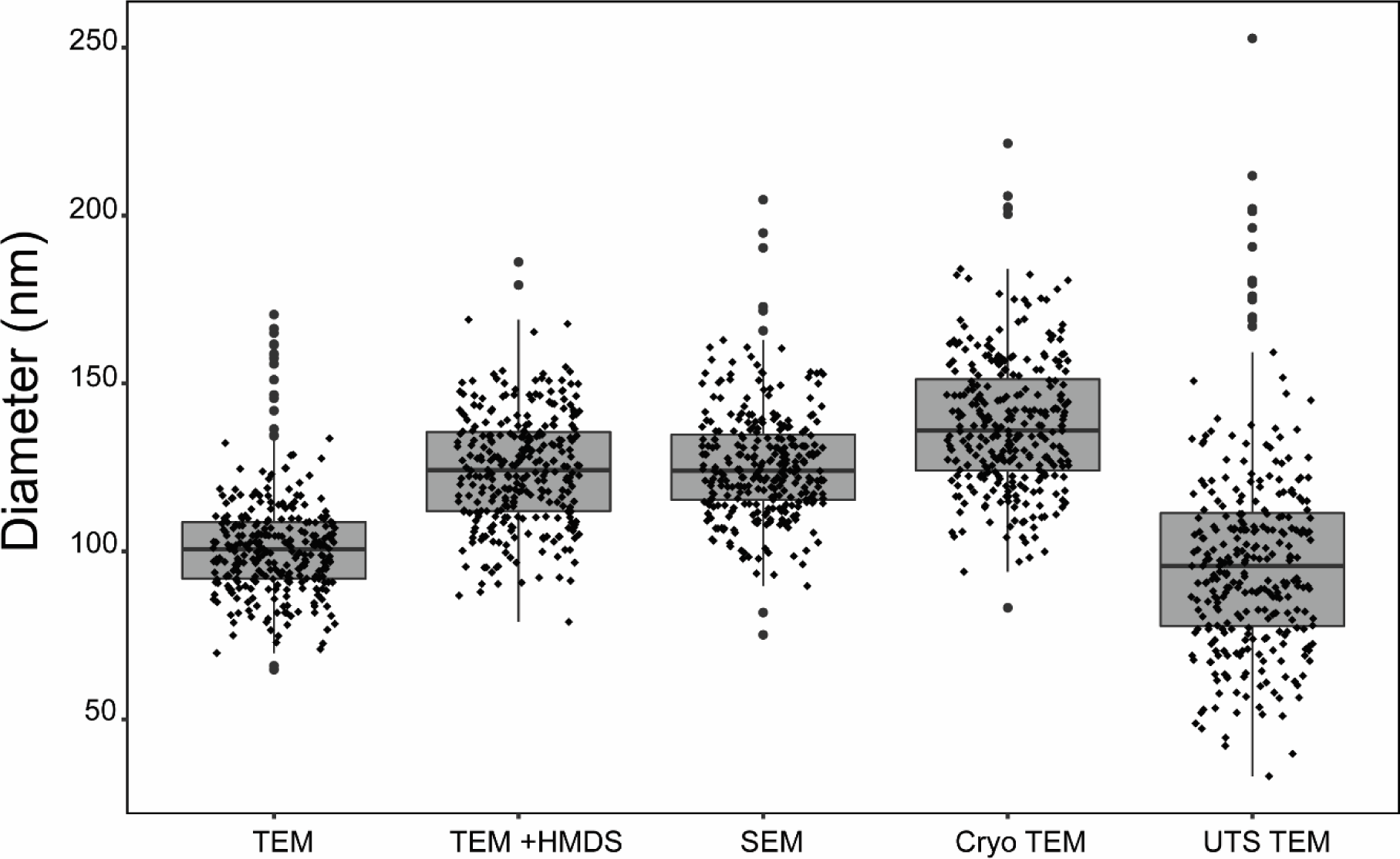
Apparent size of MCPs analyzed with different imaging techniques. Box-and-whisker plot of the size distribution of MCPs analyzed with various techniques. Note that apparent size and distribution varies widely with each technique. The only populations which are not significantly different (as defined by a p-value greater than .001 in a two-tailed t-test) are TEM of purified MCPs vs thin section TEM and SEM of purified MCPs vs TEM of dehydrated samples (p = .12 and .26, respectively). N = 300 for all, where 100 measurements were made for each of three biological replicates (three different MCP growths and purifications). Abbreviations: transmission electron microscopy (TEM), transmission electron microscopy with samples dehydrated in hexamethyldisilazane (TEM + HMDS), scanning electron microscopy (SEM), cryo transmission electron microscopy (Cryo TEM), ultra-thin section transmission electron microscopy (UTS TEM).

We attempted a number of alterations to the standard sample preparation technique to reduce MCP collapse. This included critical point drying and sample buffer exchange from the aqueous sample buffer into a high vapor-pressure solvent. These methods improved MCP structure retention, especially in samples that were exchanged into the high vapor-pressure solvent hexamethyldisilazane (HMDS) (**Fig 1B**). Overall this sample preparation technique increased the average apparent diameter of the MCPs by 22% to 124 ± 17 nm and required minimal additional steps (less than an hour of additional preparation time, even with multiple samples) (**Fig 2**). However, the exchange into HMDS led to inconsistent staining across the sample grid. This is likely due to the minimal miscibility of HMDS and the aqueous UA stain. In spite of these inconsistencies, this technique may be useful when attempting to estimate the approximate diameter of engineered MCPs using negative-stain TEM.

### Critical point drying and scanning electron microscopy reduces apparent MCP collapse

A technique that has not been widely adopted in the MCP field is scanning electron microscopy (SEM) (**Table 1**). This technique utilizes critical point drying to retain the structure of imaged samples. This is followed by treating with a sputter coater, which coats the sample in a thin layer of metal. In contrast to negative-stain TEM sample preparation, this method does not utilize an aqueous stain. For this reason, we hypothesized that critical point drying and SEM would lead to MCPs that appeared more inflated. Indeed, MCPs that were subjected to this sample preparation and imaging workflow did appear slightly more inflated than either of the negative-stain TEM methods described above (**Fig 3**). Coating for SEM also allows for tuning of the coat thickness, though there is an upper limit as increasing the metal coating thickness hindered detection of surface morphology (**Fig 3B, C**). For example, in **Fig 3B**, a coat thickness of >6 nm was used and occluded some morphological features visible in **Fig 3C**, which had a coat thickness of 6 nm. For this reason, we recommend using a minimal coat thickness (6 nm) **(Fig 3C)**, although finding a balance between optimal coat thickness, accelerating voltage, and scan speed will be necessary for each case. Overall, this technique yielded MCPs that appeared 24% (126 ± 17 nm diameter) larger, on average, than the standard negative-stain TEM method widely adopted by the field and allowed for visualization of MCP surface morphology comparable to the detail seen with negative-stain TEM (**Fig 2**). However, the additional sample preparation steps and specialized equipment may make this technique less appealing for many applications. Specifically, SEM sample preparation and imaging time per sample were approximately double that of TEM.

**Fig 3.**
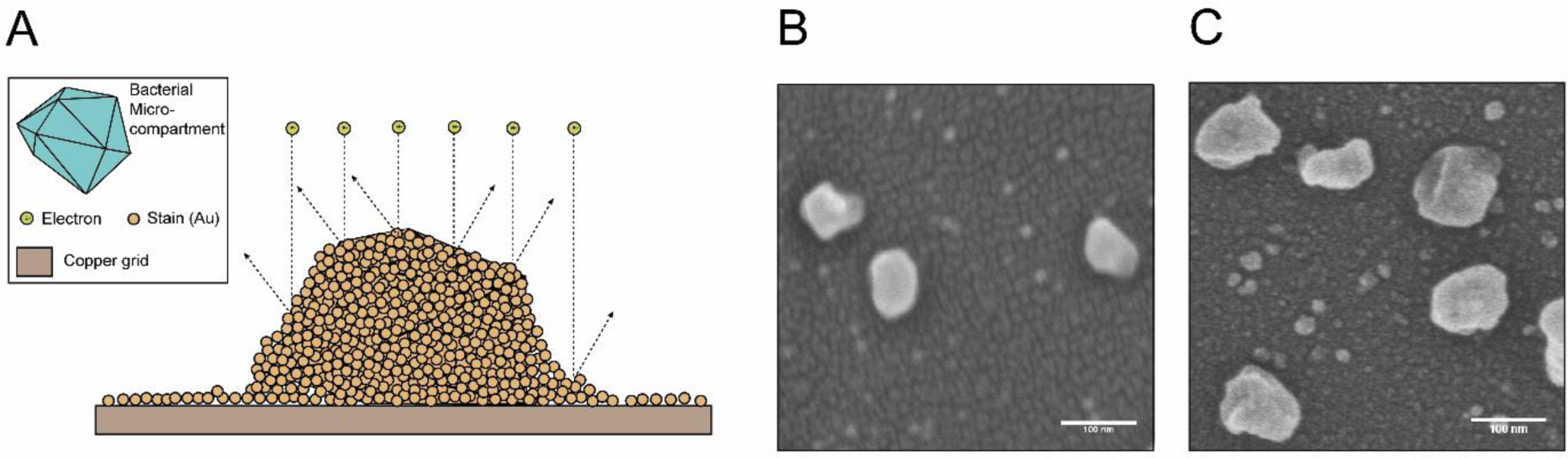
SEM of purified MCPs. (A) Schematic representation of MCPs imaged by SEM. (B) SEM of MCPs with >6 nm of gold staining. (C) SEM of MCPs with the minimal 6 nm gold coat thickness. Note that MCPs appear more inflated than in Figure 1A and surface features are apparent. Scale bars (white) are 100 nm.

### Cryo transmission electron microscopy maintains inflated MCPs

Recently, cryogenic transmission electron microscopy (cryo TEM) was used to determine the structure of an intact MCP from *Haliangium ochraceum* [56]. This MCP is unique in that it is relatively small (6.5 MDa, as opposed to the 600 MDa Pdu MCP), and regular in shape [57]. We hypothesized that because cryo TEM keeps the sample in vitreous ice and does not remove the sample from its native buffer, it would be best suited for retaining fully-inflated MCPs in their native shape and diameter (**Fig 4**). Indeed, samples that were imaged using cryo TEM produced images that on average appeared the largest of any of the techniques we attempted (138 ± 21 nm diameter). These MCPs appeared 35% larger in diameter than the standard negative-stain TEM technique and 10% larger than SEM. Samples imaged using cryo TEM also had similar variation in size observed by the other techniques, indicating that the higher average size is not due to large outliers (**Fig 2**). Indeed, cryo TEM had the second fewest outliers of any of the imaging techniques we used to assess the population size distribution.

**Fig 4.**
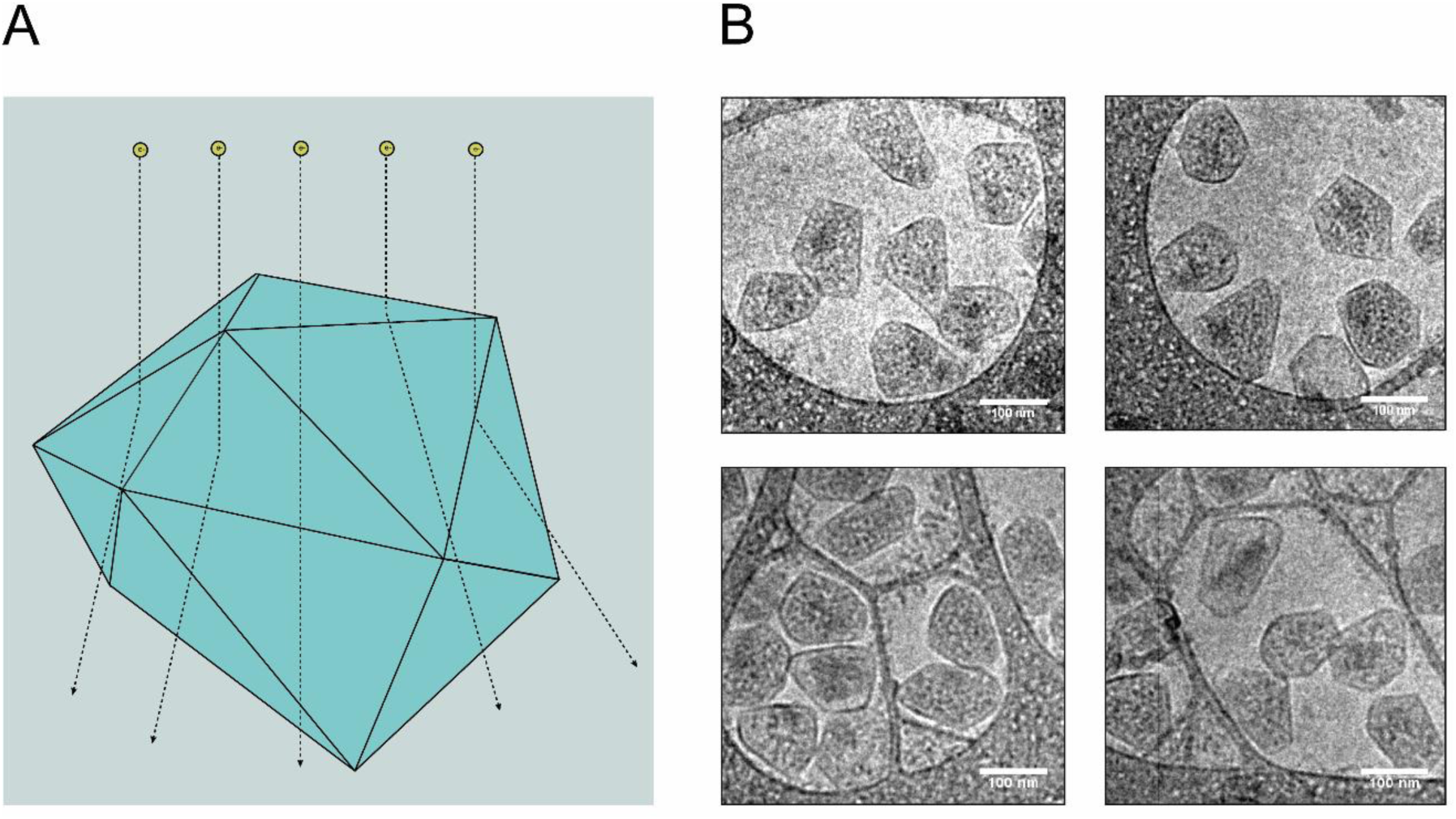
Cryo TEM of purified MCPs. (A) Schematic representation of cryo TEM of MCPs. Note that MCPs retain their native shape and are frozen in a layer of vitreous ice. (B) Micrographs of purified MCPs visualized using cryo TEM. Scale bars (white) are 100 nm.

While cryo TEM retained inflated MCPs, the lack of any contrast agent makes visualizing surface features nearly impossible. Additionally, the initial technical training for sample preparation and imaging using cryo TEM is challenging. However, an experienced microscopist can acquire cryo TEM data routinely in a single day. By contrast, chemical processing for TEM of ultra-thin sections can take several days and includes extra steps such as ultramicrotomy. Therefore, since this technique retains the native, uncollapsed state of MCPs, labs may choose to use this technique if a study is primarily focused on a change in MCP size or shape, especially on a limited number of samples (**Table 1**).

### Ultra-thin section transmission electron microscopy yields large variation in apparent size

Besides negative-stain TEM of purified MCPs, the technique most widely used in the field is TEM of ultra-thin sections. This technique has been used both on purified MCPs as well as MCPs in cells. In the earliest studies in the field, this technique was the only available option for visualizing MCPs, as a purification method was not published until relatively recently (carboxysomes were discovered in 1956 but the method for Pdu MCP purification was not published until 2012) [7,39]. This allowed for visualization of MCPs within their native context and provided researchers with a means to determine if genetic manipulations altered the expression, size, shape, and cytoplasmic distribution of MCPs.

However, TEM of ultra-thin sections has a number of drawbacks that make it a suboptimal choice for many applications. Due to the irregular shape of Pdu MCPs, ultra-thin sections produce highly variable apparent diameters (99 ± 32 nm) depending on where the MCP is sectioned (**Fig 5**). Using this technique, we found the largest variation in apparent MCP diameter, with measurements both much larger and much smaller than all previous techniques (**Fig 5**). Indeed, MCPs appeared on average 28% smaller in diameter than with cryo TEM, and the variation was between 1.5 and 1.9 times greater than all other methods (**Fig 2**). Qualitatively, MCPs visualized using TEM of ultra-thin sections appeared more rounded and less angular than with other techniques. However, this is not always the case across the field, as other labs have used this technique to produce MCP images that appear to retain their native angularity [8].

**Fig 5.**
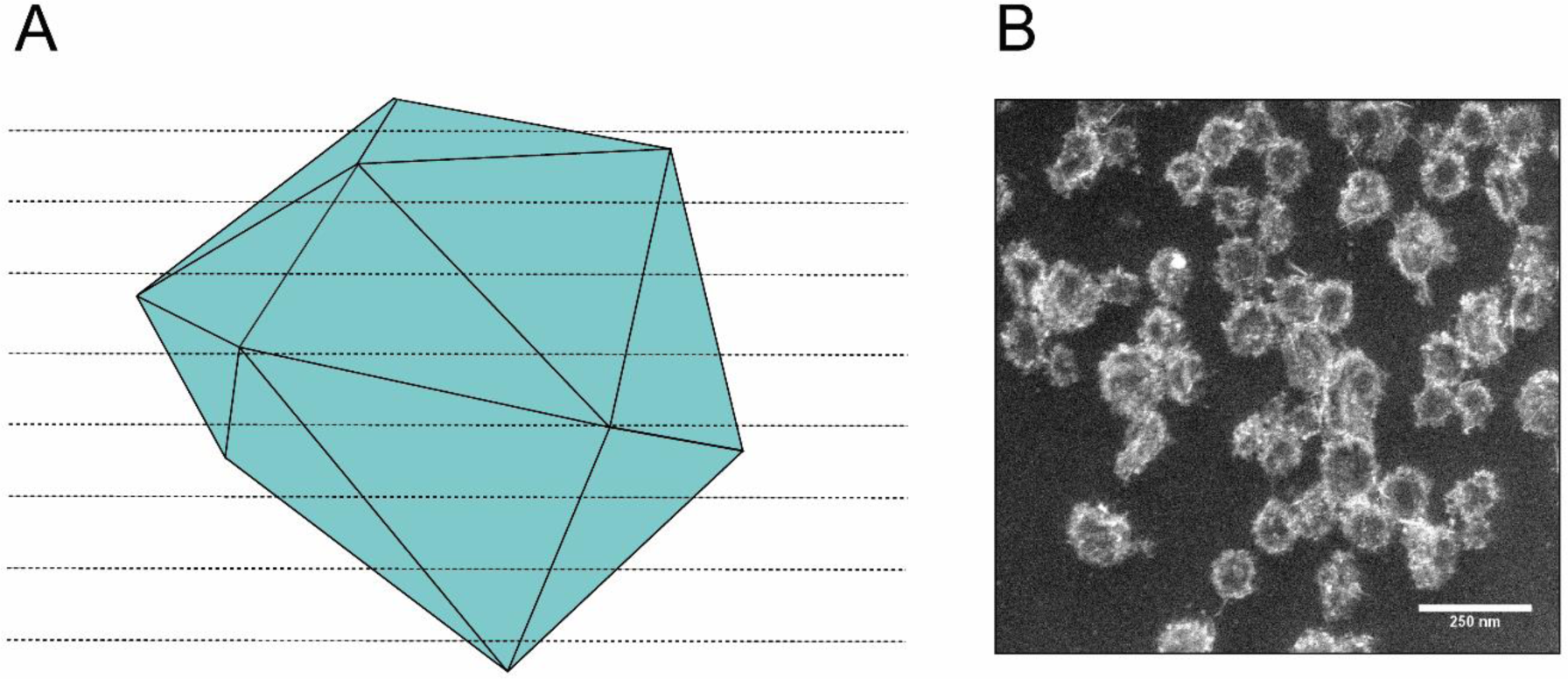
TEM of ultra-thin sections of purified MCPs. (A) Schematic representation of an MCP undergoing ultra-thin sectioning. Note that due to the irregular shape of MCPs, thin sectioning will lead to a wide range of apparent diameters. (B) Micrographs of purified and ultra-thin sectioned MCPs. Scale bar (white) is 250 nm.

Additionally, preparation of ultra-thin sections is a challenging technique to master, and it can be difficult to determine the true boundaries of MCPs when they are visualized within cells. Due to these many drawbacks, we recommend only using TEM of ultra-thin sections when it is necessary to view MCPs in their native context in the cytoplasm or when it is necessary to image the interior of MCPs (**Table 1**).

### Dynamic light scattering and nanoparticle tracking analysis enables higher-throughput MCP sizing

While microscopy allows researchers to visualize the morphology of MCPs, this may not be necessary for all studies. These imaging techniques are relatively low-throughput, and size determination is slow. One higher-throughput option for MCP sizing is particle sizing via dynamic light scattering (DLS). In this study we compared two different DLS-based techniques -- Nanosight for nanoparticle tracking analysis (NTA), and Zeta Sizer for population-based size measurements. Sizing analyses were performed on MCPs in solution (**Fig 6**), and particle size distributions (PSDs) were acquired (**Figs 6A and 6B**). When analyzed via Nanosight, the resulting distribution peak reached a maximum at 132 nm (**Fig 6A**). When analyzed via Zeta Sizer, the calculated distribution reached an intensity maximum of 122 nm (**Fig 6B**). Generally, the particle size distribution peak obtained via Nanosight was narrower than in the Zeta Sizer analysis. The average diameter measured by NTA was 149.5 ± 0.7 nm, which was larger than the 122.04 ± 0.5 nm measured by the Zeta Sizer (**Fig S2**). The disparity in mean diameter comes from large aggregates observed in the NTA experiment (**Fig S3**). To directly compare the sizing of Nanosight and Zeta Sizer, we consider differences between the mode diameter of Nanosight and the mean diameter (Z_ave_) of the Zeta Sizer to be the most accurate comparison. The mode diameter of 130.7 ± 1.0 nm is slightly higher than the measured 122.04 ± 0.5 nm observed in Zeta Sizer measurements. Finally, the polydispersity index (PDI) calculated via Zeta Sizer was 0.045 ± 0.001, indicating that purified MCPs are monodisperse, as expected. We attribute discrepancies in diameter measurements to differences between the measurement techniques and their respective calculations of particle diameter. The full experiment report obtained for NTA measurements is shown in **S3 Fig.**

**Fig 6.**
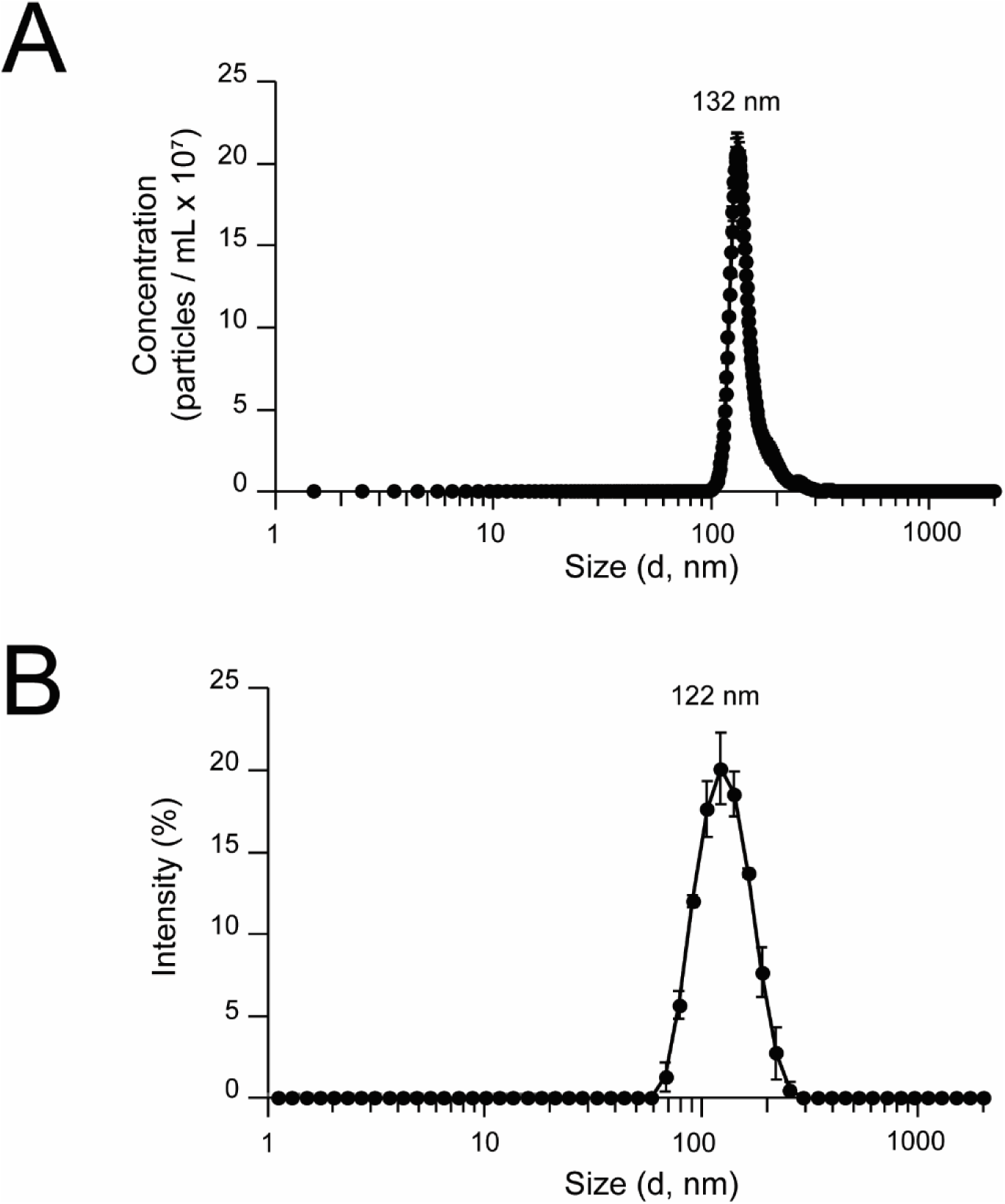
Higher-throughput sizing of purified MCPs using DLS. Sizing MCPs in solution via light scattering techniques. (A) Particle size distributions measured via Nanosight, and (B) Zeta Sizer.

To further assess the stability of DLS sizing measurements over an order of magnitude of concentrations of MCPs, we compared size measurements at 50 µg/mL and 500 µg/mL MCPs using Zeta Sizer (**S4 Fig**). Importantly, the raw correlation data obtained at 50 µg/mL and 500 µg/mL were in good agreement; the resulting Z_ave_ values calculated were 121.1 ± 0.32 nm and 122.0 ± 0.51 nm, respectively **(S4A and S4B Figs)**. Polydispersity indices obtained for MCPs at 50 µg/mL and 500 µg/mL were 0.069 ± 0.001 and 0.045 ± 0.001, respectively, indicating a high degree of uniformity of MCPs (**S4C Fig**). Full intensity, number, and volume PSDs for DLS measurements are presented in **S4D-F Figs.** As expected, we observed similar PSDs for measurements collected at 50 µg/mL and 500 µg/mL MCPs. Intensity PSDs displayed maximum intensities at ∼122 nm. Number and volume PSDs displayed maxima near ∼90 nm, and were left-shifted with respect to the intensity PSDs. Slight shifting of the PSDs between intensity, number, and volume distributions is most likely due to the non-spherical nature of MCPs. The consistency and stability of DLS measurements over an order of magnitude of concentration indicate that Zeta Sizer is a suitable technique for analysis of MCPs over a range of concentrations. Notably, the diameters obtained by Zeta Sizer appear more similar to the results obtained by SEM or TEM samples treated with HMDS but are 12-13% smaller than MCPs observed by cryo TEM. However, Nanosight results appeared most similar to those obtained by cryo TEM (132 nm vs. 138 nm).

## Conclusion

Our results suggest that the technique used to visualize and measure MCPs can alter how we interpret our experimental results. This is especially important when comparing results across studies which used different techniques to assess their results. Our hope is that this study can provide a guideline for the appropriate use of each of the many available techniques used in the field to assess MCPs. Our results can also be used to contextualize and compare results across different studies by providing approximate percent changes in apparent size for each technique.

## Acknowledgements

The authors would like to thank Robert Colby, Mark Heinnickel, and Giovanni Pilloni for their helpful thoughts on additional methods to include in this work, and members of the Tullman-Ercek lab for helpful comments during the preparation of this manuscript.

## Supporting information

**S1 Table.** Reported sizes of MCPs. The reported size range for MCPs, the work in which the size was reported, the system being analyzed (Propanediol utilization (Pdu), Ethanolamine utilization (Eut), Carboxysome (Carb)), and the technique used for the analysis (TEM of ultra-thin sections (TEM UTS), TEM of purified MCPs (TEM Pur)).

| **Size** | **Report** | **Type** | **Method** |
| --- | --- | --- | --- |
| 77-123 nm | Parsons, JBC, 2008 | Pdu | TEM UTS |
| 100-150 nm | Havemann, Journal of Bacteriol, 2003 | Pdu | TEM Pur |
| 88-106 nm | Schmid, JMB, 2006 | Carb | Cryo EM |
| 100 nm | Shively, Journal of Bacteriol, 1970 | Carb | TEM UTS |
| 100-200 nm | Bobik, Journal of Bacteriol, 1999 | Pdu | TEM UTS |
| 100-200 nm | Choudhary, PLoS ONE, 2012 | Eut | TEM UTS |
| 110-220 nm | Slininger Lee, ACS Syn Biol, 2017 | Pdu | TEM Pur |
| 100-150 nm | Chowdhury, MMBR, 2014 (no citation - unpublished Bobik) | Pdu | none |

**S1 Fig.**
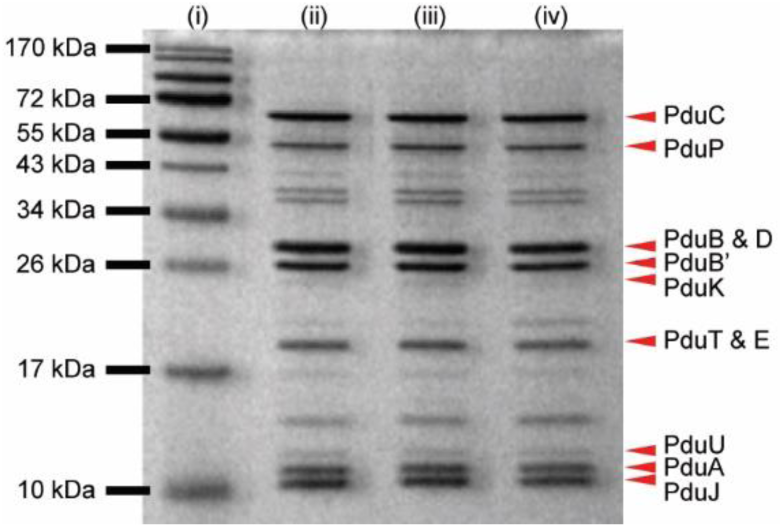
Coomassie-stained gel of three biological replicates used for EM images. Lanes: (i) molecular weight standard, (ii-iv) replicates of purified Pdu MCPs.

**S2. Fig.**
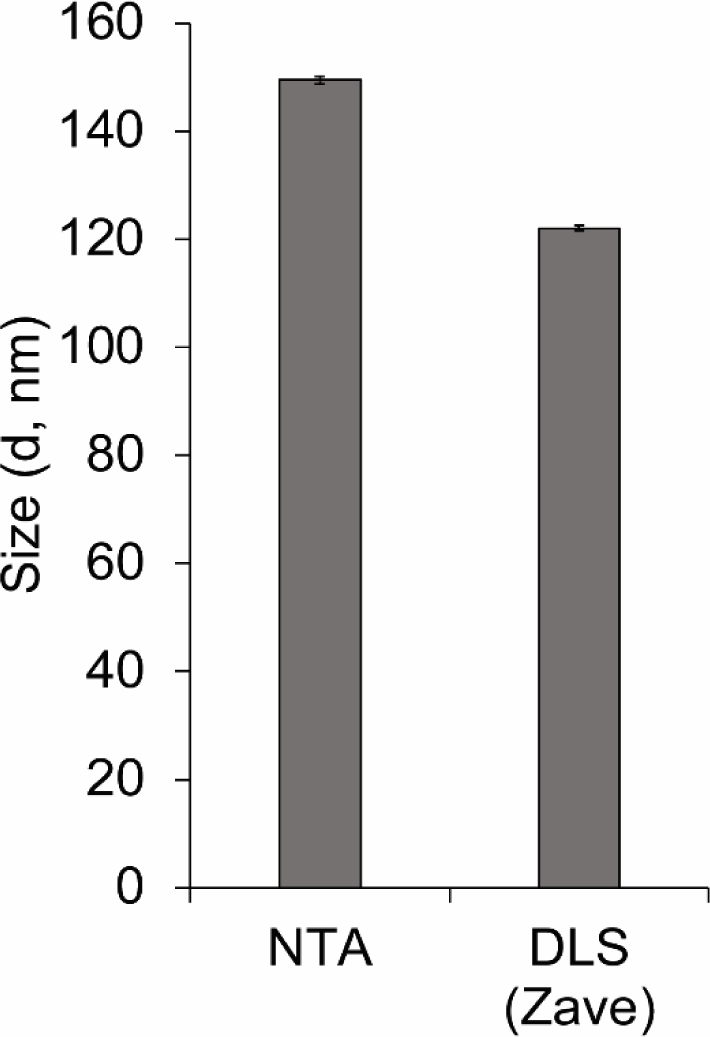
Comparison of average diameters measured via NTA and DLS. Error bars represent the standard deviation.

**S3 Fig.**
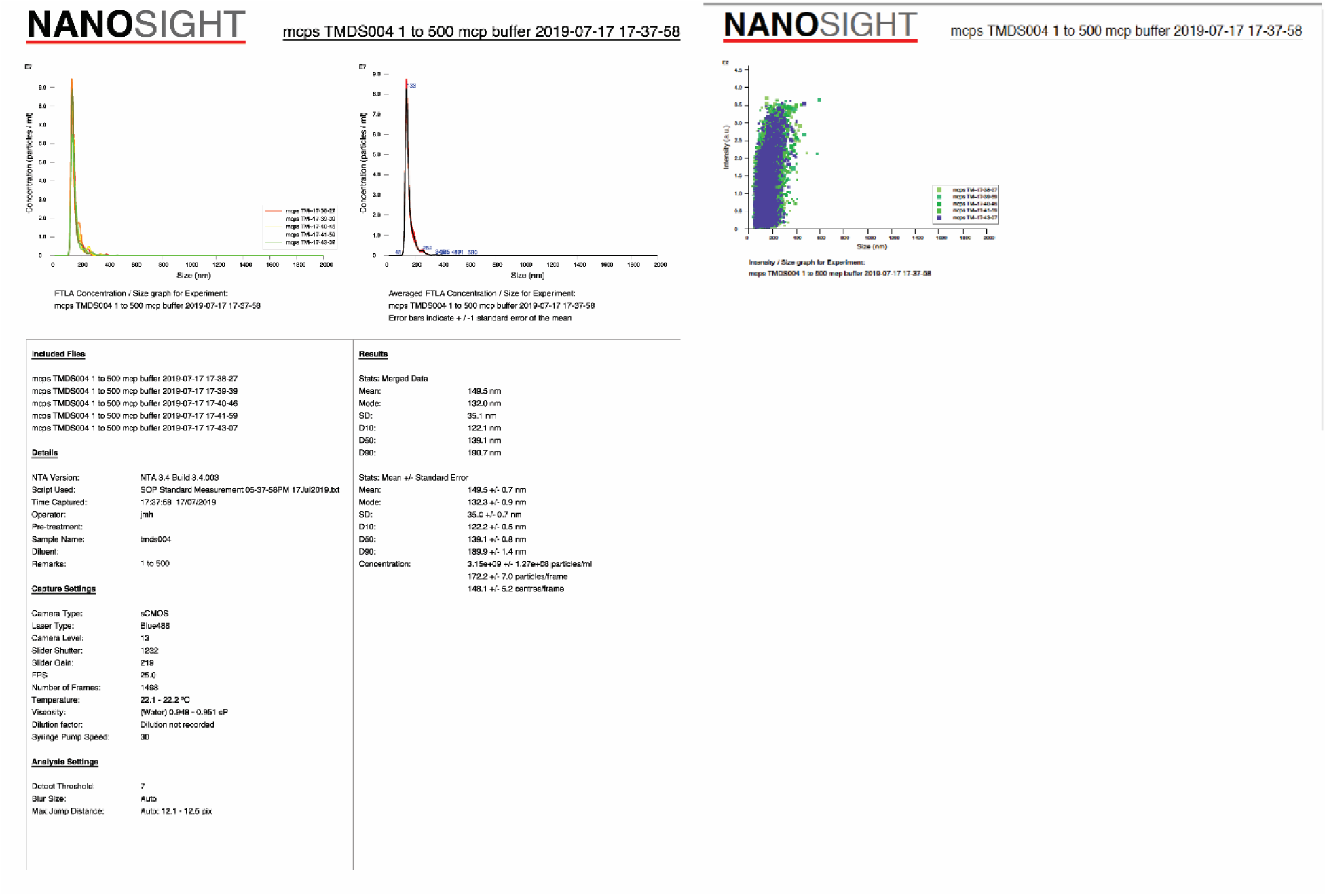
Full Nanosight/NTA analysis report used in this study.

**S4 Fig.**
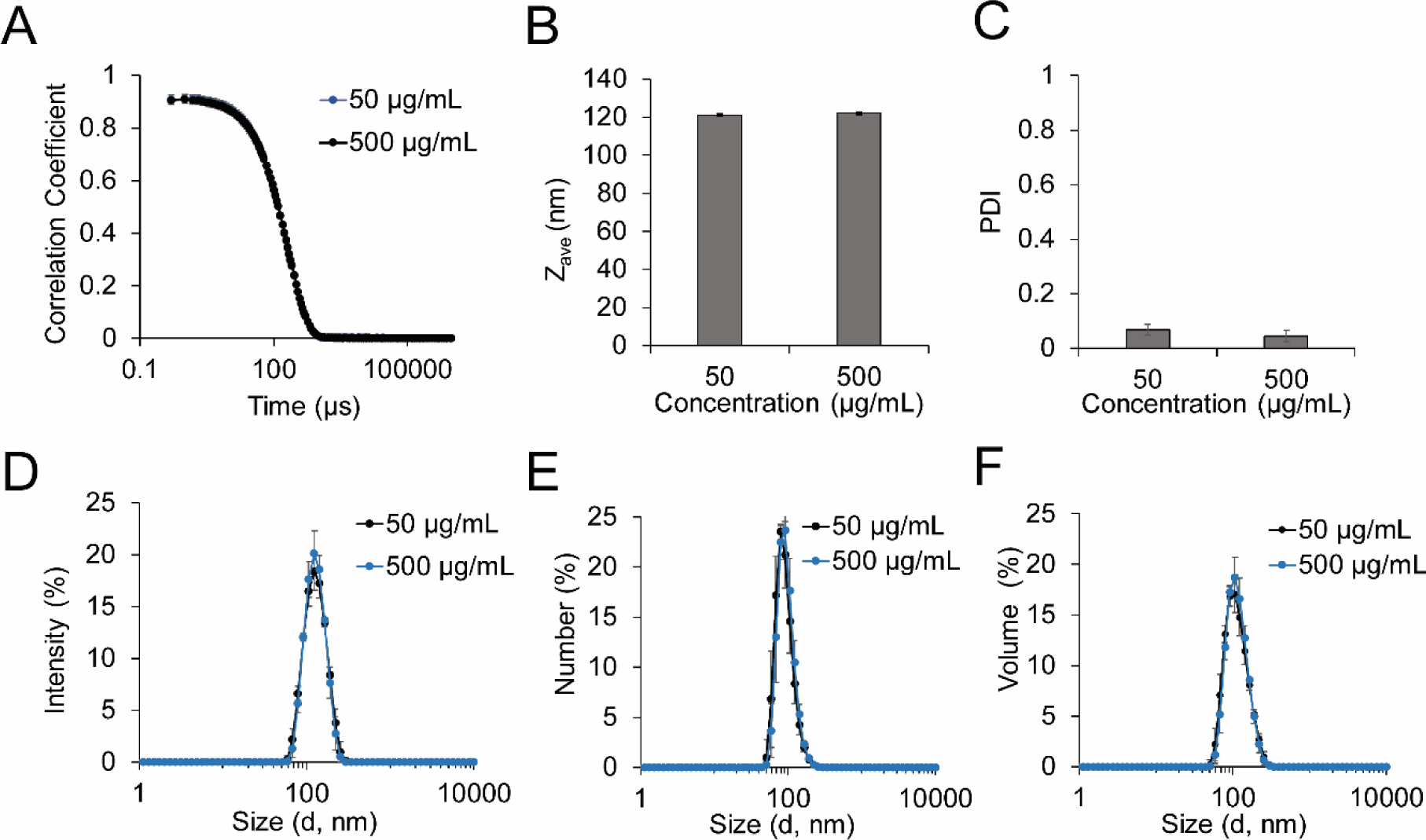
DLS analysis of MCPs at 50 µg/mL and 500 µg/mL. Raw correlation data (A), calculated Zave (B), and polydispersity indices (PDI) (C) of MCPs. Intensity (D), number (E), volume (F) particle size distributions of MCPs.

**S1 File. Raw sizing data from images.**

**S2 File. Raw sizing data from Zeta Sizer.**

**S3 File. Raw sizing data from Nanosight.**

**S4 File. Raw, uncropped SDS-PAGE gels of purified MCP samples.**

