## Supplementary material for "Apparent size and morphology of bacterial microcompartments varies with technique": S4_File

**Technical Replicate 1 20181016**

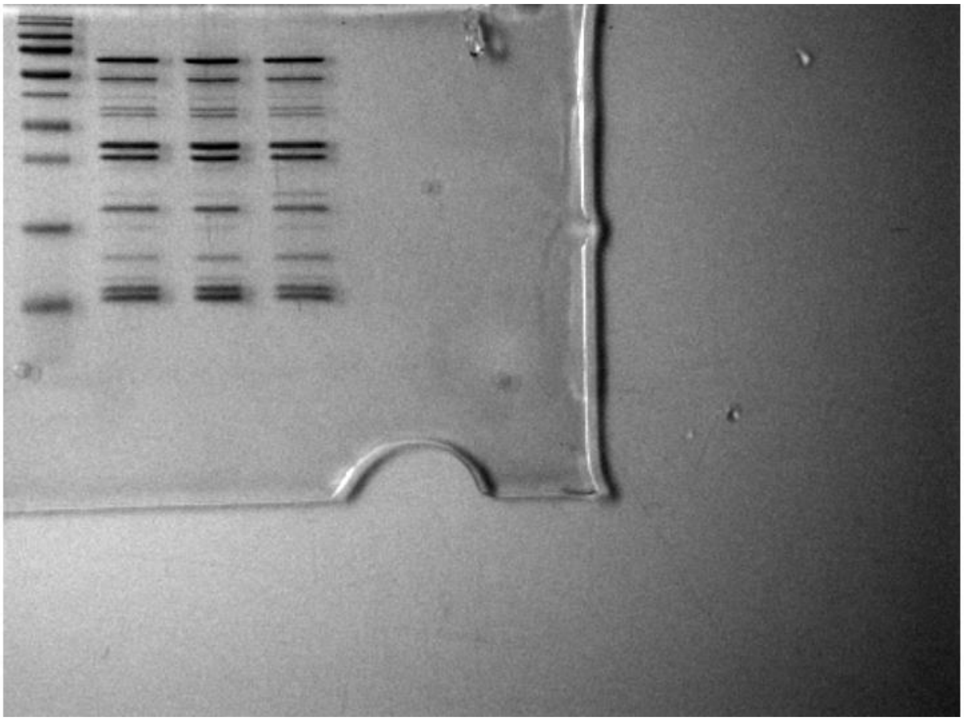

**Technical Replicate 2 20181016**

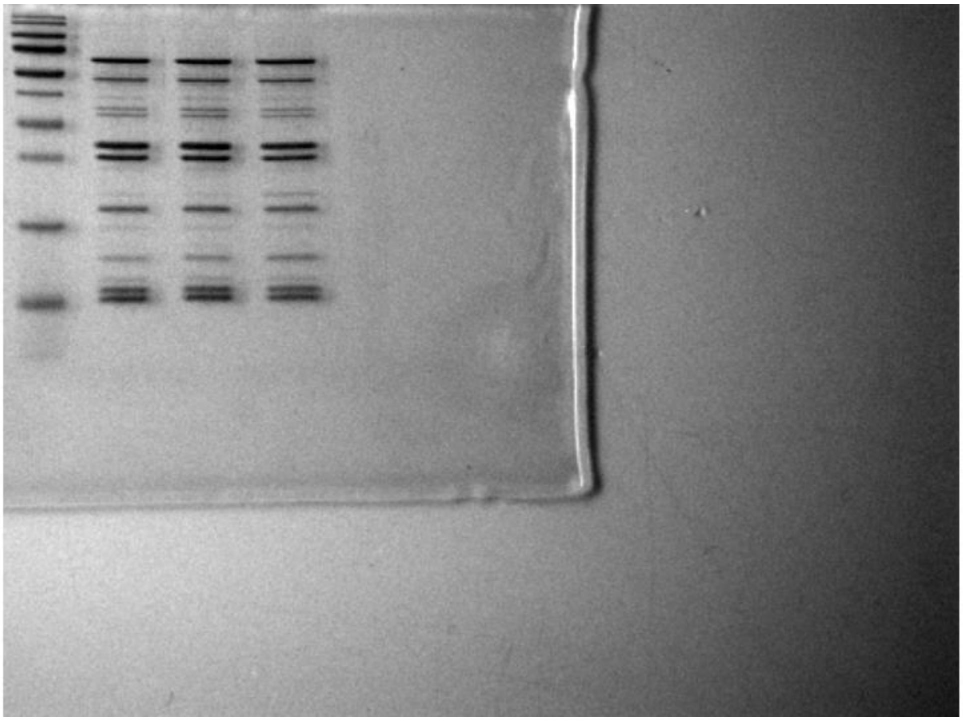

**Technical Replicate 1 20181113**

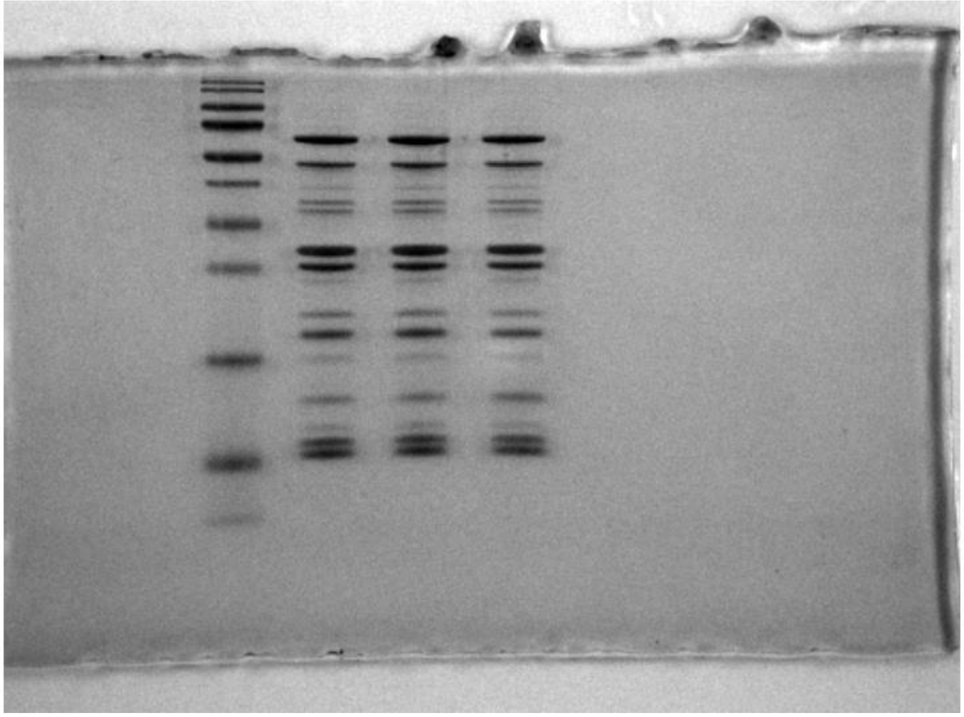

**Technical Replicate 2 20181116**

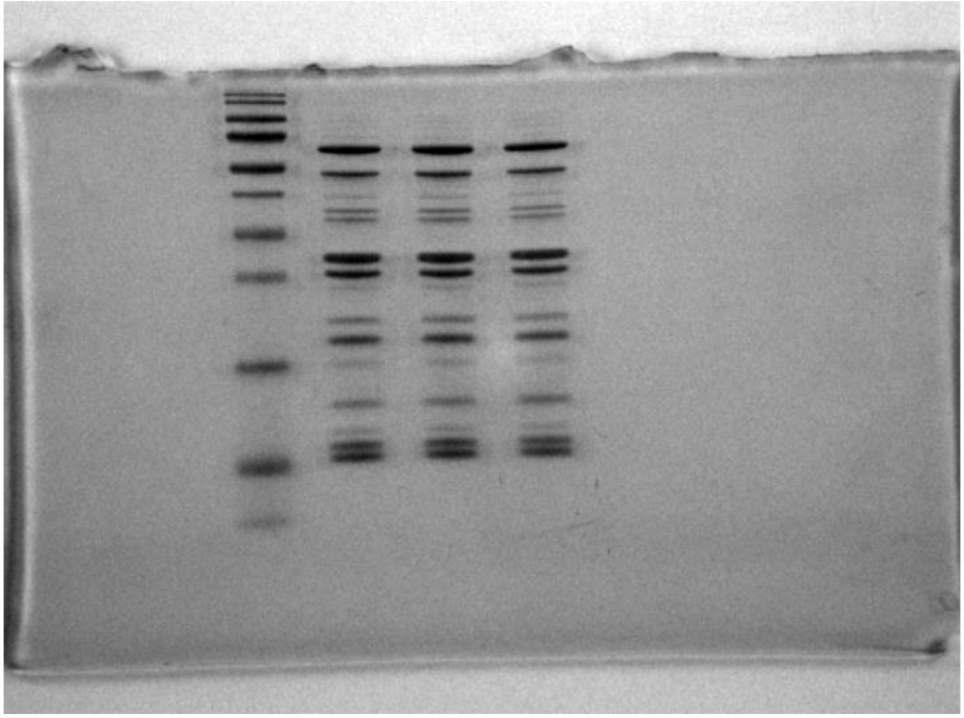

**Coomassie 20190208**

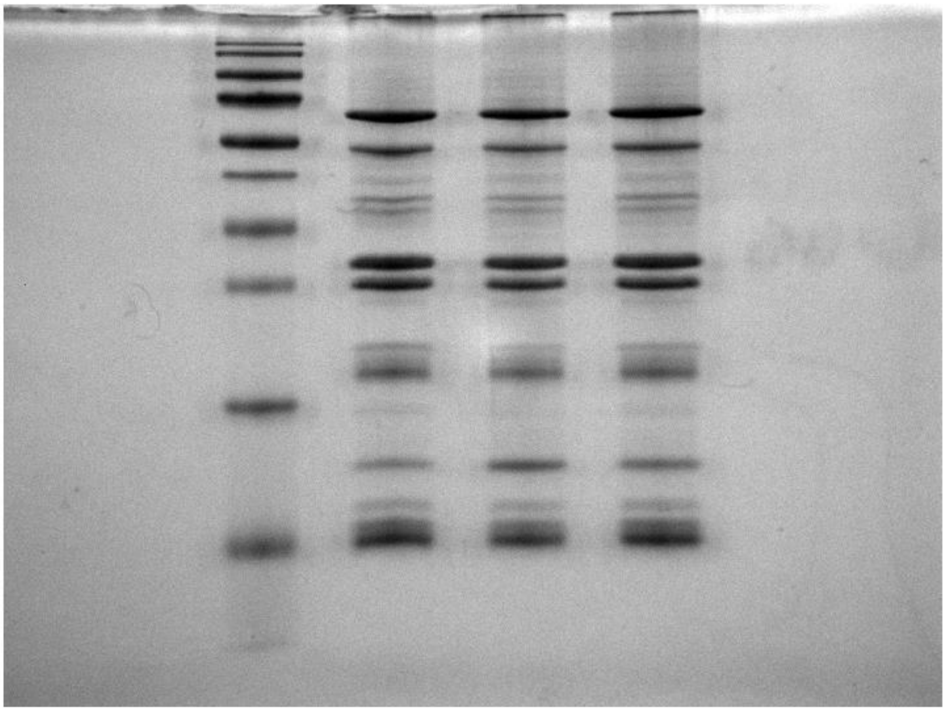
